# Actin shells control buckling and wrinkling of biomembranes

**DOI:** 10.1101/781153

**Authors:** Remy Kusters, Camille Simon, Rogério Lopes Dos Santos, Valentina Caorsi, Sangsong Wu, Jean-Francois Joanny, Pierre Sens, Cecile Sykes

## Abstract

Global changes of cell shape under mechanical or osmotic external stresses are mostly controlled by the mechanics of the cortical actin cytoskeleton underlying the cell membrane. Some aspects of this process can be recapitulated *in vitro* on reconstituted actin-and-membrane systems. In this paper, we investigate how the mechanical properties of a branched actin network shell, polymerized at the surface of a liposome, control membrane shape when the volume is reduced. We observe a variety of membrane shapes depending on the actin thickness. Thin shells undergo buckling, characterized by a cup-shape deformation of the membrane that coincides with the one of the actin network. Thick shells produce membrane wrinkles, but do not deform their outer layer. For intermediate micrometer-thick shells, wrinkling of the membrane is observed, and the actin layer is slightly deformed. Confronting our experimental results with a theoretical description, we determine the transition between buckling and wrinkling depending on the thickness of the actin shell and the size of the liposome. We thus unveil the generic mechanism by which biomembranes are able to accommodate their shape against mechanical compression, through thickness adaptation of their cortical cytoskeleton.

## Introduction

Cellular deformations are produced by a wealth of active dynamical processes occuring near the cell membrane, locally curving the membrane inwards or outwards. Most cells display a cytoskeleton underneath their lipidic membrane. Such composite objects are able to adapt to various external stresses by changing their shape and mechanics, and therefore optimize their movement within the body. The “simplest” example of a membrane-cytoskeleton cell is the red blood cell (RBC), with its thin spectrin cytoskeleton. RBC shapes can vary among discocyte, stomatocytes and echinocytes by possible changes of spontaneous curvature of the membrane (1). Different RBC shapes display distinct mechanical behaviors, as revealed by an analysis of their temporal undulations (1, 2).

The spherical volume enclosed by a thin elastic shell can be reduced, for example, by an osmotic shock. In that case, compressive stresses build up that cause the formation of sharp points and bends to develop. An initial spherical shell can undergo “buckling”, and a bowl- or cup-like shape is reached if the bend is localized. In the case of multiple sharp points that form at the same time, the final shape appears wrinkled or crumpled. Buckling, crumpling and wrinkling by volume reduction are therefore unambiguous signatures of the solid elastic properties of the shell material. Exact buckled shapes can be described by an energy optimization of bending and stretching (3–5) and such a mechanism applies to thin polymer shells or fluid droplets coated with solid colloidal particles (Pickering emulsions) (6–8). The obtained cup-like shapes are reminiscent of stomatocytes and provide a conceptually simple way to produce colloids with two different sides (Janus colloids) (6). Buckling of thin elastic polymer shells occurs under a critical osmotic shock that increases when the shell is thicker (9).

Cancer cells are sensitive to their external mechanical environment and, under stress, activate changes in their cytoskeleton (10). Conversely, the cytoskeleton, and in particular the cell cortex, the thin actin network underneath the cell membrane, can monitor cell shape changes (11). In all cases, global shape changes are directly impacted by the mechanics of the cytoskeleton and small perturbations of the cortex structure may deeply change the equilibrium shape and mechanical properties of the cells. Examples are alterations in the cross-linked spectrin network of red blood cells (12) or perturbations of the cortical actin layer of eukaryotic cells (13).

Here, we use a model experimental system of a liposome coated on its external surface by a dynamic actin layer organized in a branched filamentous network. This system mimics the cell cortex but in the absence of any myosin molecular motor. Our experimental setup allows for a fine control of the branched, polymerized, actin layer. Under spontaneous evaporation of the solution, water leaks out of liposomes and we observe both buckling and wrinkling under spontaneous volume loss. A controlled change of liposome volume is applied through a fixed hyper-osmotic shock which led us to observe a buckling of the actin network layer that matches the shape of the liposome membrane. We interpret our experimental observations in light of theoretical developments in the context of thin polymer shell buckling.

We further study how the shape of the liposome-shell system changes upon a volume decrease, when the thickness of the grown actin layer is varied. Wrinkling is favored for thicker shells over buckling. In cells with an underlying cytoskeleton, the thickness of the cytoskeletal network is therefore a way of protecting from, or providing, buckling.

## Methods

### Reagents, lipids and proteins

Chemicals are purchased from Sigma Aldrich (St. Louis, MO, USA) unless specified otherwise. L-alpha-phosphatidylcholine (EPC), 1,2-distearoyl-sn-glycero-3-phosphoethanolamine-N-[biotinyl polyethylene glycol 2000] (biotinylated lipids). TexasRed ^©^ 1,2-dipalmitoyl-sn-glycero-3-phosphocholine, triethylammonium salt is from Thermofisher (Waltham, Massachussetts, USA). Actin is purchased from Cytoskeleton (Denver, USA) and used with no further purification. Fluorescent Alexa Fluor 488 actin conjugate is obtained from Molecular Probes (Eugene, Oregon, USA). Porcine Arp2/3 complex is purchased from Cytoskeleton and used with no further purification. Mouse *α*1*β*2 capping protein (CP) is purified as in (14). Untagged human profilin and Streptavidin-pVCA (S-pVCA, where pVCA is the proline rich domain-verprolin homology-central-acidic sequence from human WASP, starting at amino acid Gln150) are purified as in (15). A solution of 30 *μ*M monomeric actin containing 15% of labeled Alexa Fluor 488 actin conjugate is obtained by incubating the actin solution in G-Buffer (2 mM Tris, 0.2 mM CaCl_2_, 2 mM ATP, 0.2 mM DTT, pH 8.0) overnight at 4°C. The isotonic (resp. hypertonic) working buffer contains 95 mM (resp. 320 mM) sucrose, 1 mM Tris, 50 mM KCl, 2 mM MgCl2, 0.1 mM DTT, 2 mM ATP, 0.02 mg/mL *β*-casein, adjusted to pH 7.4. Osmolarities of the isotonic and hypertonic working buffers are respectively 200 and 400 mOsmol, as measured with a Vapor Pressure Osmometer (VAPRO 5600). All proteins (S-pVCA, profilin, CP, actin) are mixed in the isotonic working buffer.

### Preparation of liposomes with actin cortices

Liposomes are prepared using the electroformation technique (16). Briefly, 10 *μ*L of a mixture of EPC lipids, 0.1% biotinylated lipids and 0.5% TexasRed lipids with a concentration of 2.5 mg/ml in chloroform/methanol 5:3 (v/v) are spread onto indium tin oxide (ITO)-coated plates under vacuum for 2 h. A chamber is formed using the ITO plates (their conductive sides facing each other) filled with a sucrose buffer (0.2 M sucrose, 2 mM Tris adjusted at pH 7.4) and sealed with hematocrit paste (Vitrex Medical, Denmark). Liposomes are formed by applying an alternating current voltage (10 Hz, 1 V rms) for 2 h.

Liposomes are incubated for 15 min with 350 nM of S-pVCA, the activator of the Arp2/3 complex, which binds to the biotinylated lipids of the membrane through the streptavidin tag. Actin polymerization starts instantaneously when S-pVCA-coated liposomes are placed in a mix containing a final concentration of 3 *μ*M monomeric actin (15% fluorescently labeled with Alexa Fluor 488), 3 *μ*M profilin, 37 nM Arp2/3 complex and 25 nM CP. Different thicknesses of actin shells are obtained depending on the polymerization time, set from 5 to 30 min.

### Osmotic shock on liposomes after actin polymerization

The actin polymerization reaction is stopped by diluting liposomes twice either in isotonic working buffer (200 mOsmol, control) or in hypertonic working buffer (400 mOsmol, osmotic shock) and incubated for 15 min. After the osmotic shock, the final solution is at 300 mOsmol resulting in a 100 mOsmol osmotic pressure difference.

### Image acquisition of liposomes and actin shells

Epifluorescence (GFP: LED with wavelength 460nm (CoolLED pE-4000), emission filter: ET542/20x; Texas red: LED with wavelength 580nm (CoolLED pE-4000), emission filter: ET610LP) and phase contrast microscopy are performed using an IX70 Olympus inverted microscope with a 100x oil-immersion objective. Images are acquired by a charge coupled device CCD camera (CoolSnap, Photometrics, Roper Scientific).

### Liposome morphology and actin shell analyses

Image analyses are performed with ImageJ software and data are processed on Matlab and Python. The position of the liposome surface is defined by the peak of the membrane fluorescence profile. The actin network thickness, *h*_0_, is defined as the distance between the position of the membrane and the one of the half maximum amplitude of the actin fluorescence profile. Liposome shapes are automatically tracked with the ImageJ plugin: JFilament 2D, where the tracking parameters “beta”, “deform iterations” and “weight” are respectively set to 6000-10000, 50 and 10.5. The shape is manually segmented for the plugin to track boundaries. From these tracks, we evaluate the curvature at each point along the curvilinear abscissa, *s*, using the method of interpolating splines. We then infer the maximal curvature, *C_max_*. From the tracks, we also evaluate the mean radius of the deformed liposome *R*. We suppose that the liposome area is constant and we use the method of least squares, i.e. fitting a circle that minimizes the square deviation. We estimate the un-deformed, reference, radius of the liposome as 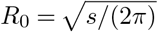. Once *R* and *R*_0_ are determined, we estimate the relative reduction of volume due to the osmotic shock 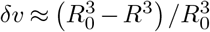. Note that this approximation is valid for small values of *δυ*. To classify the shapes into either buckled or wrinkled, 93 vesicles were analyzed and manually categorized ^1^. Wavelengths of wrinkled shapes are obtained by applying a Fourier transform to the shape radius as a function of angle (examples in Supplementary Figure). The chosen wavelength is then the one with highest spectral energy.

## Results and discussion

### Buckling and wrinkling of actin shells

The growth of a branched actin network from the surface of liposomes forms a spherical actin shell that thickens over time, as previously characterized (Fig. 1A and (15)). Whereas stresses may develop within the actin shell and eventually break the shell open and develop a comet for motility (15, 17), here, we focus our observations on homogeneous actin shells. We observe liposomes before they undergo a symmetry breaking event of the actin network. A typical liposome covered with an actin shell made of a branched network is given in Fig. 1A. When the preparation evaporates spontaneously, we observe that liposomes covered with an actin shell tend to deform irreversibly either into a “buckled” shape or a “wrinkled” shape, and this happens progressively within hours (Fig. 1B, top). In the absence of actin, liposome deformations differ drastically: thermal fluctuations varying in space and time appear around the spherical shape of the liposome that never irreversibly deforms (Fig. 1B, bottom and Supplementary movie 1). The latter case is well known and monitored by the bending energy (or bending rigidity) of lipid membranes, of the order of a few *kT*, close to thermal energy (18). Here, evaporation provokes an osmotic unbalance by increasing solute concentration outside the liposomes, water leaks out to restore the osmotic balance and the liposome volume decreases. As a consequence, liposome membrane tension decreases and therefore the amplitude of thermal fluctuations increases. As soon as actin covers liposomes, thermal fluctuations are not visible. Instead, buckled and wrinkled shapes are observed that continue to develop when spontaneous evaporation is maintained. In order to control the volume change of liposomes, we now apply a quantified osmolarity change to the external solution.

**Fig. 1.**
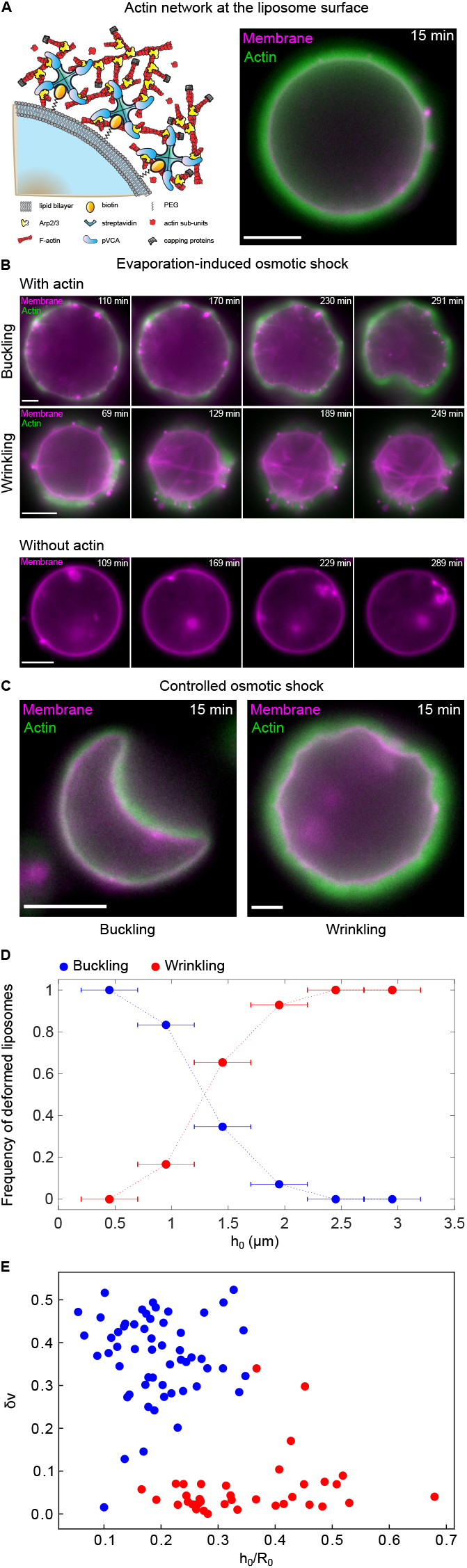
A. (Left) Scheme of the experimental system; proteins not to scale. (Right) Merged example of membrane (magenta) and actin network (green) under isotonic conditions. Scale bars, 5 *μ*m. B. (Top) Time lapses of membrane buckling and wrinkling induced by evaporation-based osmotic shock, at room temperature, on liposome covered with an actin shell. Time indicated is the time after actin polymerization stops. (Bottom) Time lapse of membrane deflation induced by evaporation-based osmotic shock on a naked liposome. C. Buckling and wrinkling induced by controlled osmotic shock on liposome covered with an actin shell. Epifluorescence images of membrane (magenta) and actin (green). Scale bars, 5 *μ*m. D. Frequency of liposomes displaying buckling or wrinkling as function of the actin thickness *h*_0_. Segmented lines are guides to the eyes. E. Wrinkled (red) and buckled (blue) shapes displayed as a function of volume change *δυ* and the ratio of the actin thickness *h*_0_ over the liposome radius *R*_0_.

### Effect of an osmotic shock on liposomes coated with an actin shell

We apply a controlled osmotic shock of 100 mOsm (corresponding to 25 kPa) by diluting twice the liposomes surrounded by an actin shell (solution at 200 mOsm) in a hypertonic solution (400 mOsm). After the osmotic shock, we obtain the same two types of shapes as observed under spontaneous evaporation: either a cup or a wrinkled shape (Fig. 1C). Cup shapes are reminiscent of the ones of buckled spherical capsules under an osmotic shock (5, 9) and the wrinkled shapes are reminiscent of that of a liposome under acto-myosin contraction (19, 20). Again, we confirm that naked liposomes subjected to a decrease in volume (osmotic shock) display dynamical thermal shape undulations (Supplementary movie 2). In contrast, actin-covered liposomes do not display visible thermal shape fluctuations, but instead, once deformed, their shapes remain over time. Their morphological response to the osmotic shock is a signature of the elasticity of the actin shell surrounding the liposome and the thickness of the elastic shell determines its capacity to undergo buckling (9). Our experimental system allows for a perfect control of actin network thickness by varying the actin polymerization time, which monitors the quantity of actin that polymerizes within the growing network (17). We obtain an actin shell thickness, *h*_0_, ranging from 250 nm to 3 *μ*m. We observe that liposomes with a thin actin shell are more likely to undergo buckling under an osmotic shock, whereas wrinkling is favored for thick actin shells (Fig. 1D). We represent the occurrence of buckling and wrinkling for all the analysed liposome shapes after osmotic shock in the 2-dimensional space (*h*_0_/*R*_0_, *δυ*) in Fig. 1E. We find indeed a clear separation between the two families of shapes (buckling and wrinkling). To rationalize these findings, we study theoretically the two types of deformations.

### Buckled shapes

We consider an initially spherical liposome with a radius *R*_0_ and an area 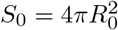 before the osmotic shock. Due to the osmotic compression, the mean radius of the deflated liposome decreases to *R* and the relative decrease of its inner volume is 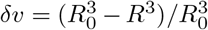 or *R* ≃ *R*_0_(1 – *δυ*/3), to first order in *δυ*. Buckling is characterized by a cup shape with a large concave region (Fig. 1C). The buckling of elastic shells is a well known phenomenon (3, 4), and can be understood analytically as follows (3). For moderate buckling, the buckled shape remains axi-symmetric and the deformed (concave) region is almost the mirror image of its initial, non-deformed shape, with most of the elastic energy concentrated in a narrow “bending strip” of width *a* connecting the convex and concave regions (Fig. 2B). We call *r* and *d* the radius and depth of the concave region respectively and *α* is the polar angle defining the latitude of the bending strip (Fig. 2B). Assuming *r* ≪ *R*_0_ gives the following geometrical relations: *r*/*R*_0_ = sin *α* ~ *α* and *d*/*R*_0_ = 2(1–cos*α*) ~ *α*^2^. The relative volume decrease is expressed as 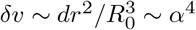. The width *a* of the bending strip is then obtained by balancing bending and stretching stresses in the bending strip. Calling *ζ* the typical radial displacement of the shell in the bending strip, the curvature, which is locally maximal, is of order *C_max_* ~ *ζ*/*a*^2^. An actin network is slightly compressible (21), but we consider it here as incompressible for sake of simplicity (Poisson coefficient equal to 1/2). For thin shells of elastic Young modulus *E* and thickness *h*_0_, we obtain the bending energy density 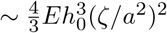. Compression in the bending strip occurs primarily along circles of constant latitude with a strain tensor of order *ζ*/*R*, giving a stretching energy density ~2*Eh*_0_(*ζ*/*R*_0_)^2^ in agreement with (3). Moreover, matching the edges of the bending strip with the convex and concave regions of the buckled shell imposes that *C_max_* ~ *α*/*a*, or *ζ* ~ *αa*. Taking this constraint into account and integrating the energy densities over the area of the bending strip ~ *ra* yields the total buckling energy,

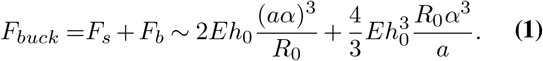

**Fig. 2.**
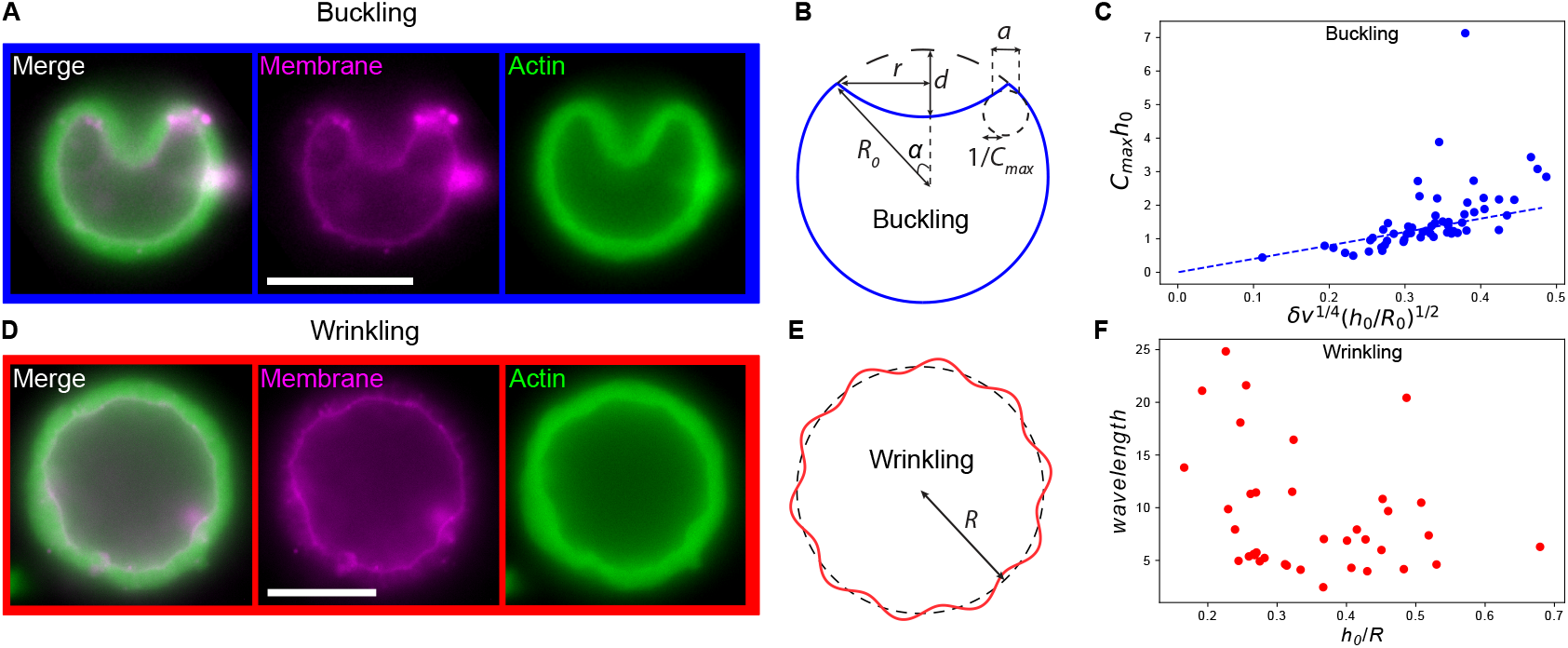
Analysis of buckled and wrinkled shapes. Example epifluorescence images of buckling (A) and wrinkling (D) with membrane (magenta) and actin (green). Scale bars, 5 *μ*m. Schematic representation of buckling (B) and wrinkling (E) induced by controlled osmotic shock, with shape parameters. C. Buckled shapes, maximal radius of curvature multiplied with the shell thickness, *C_max_h*_0_ as function of the osmotic compression *δυ*. The dashed line represents the analytic prediction of Eq. 3, with a proportionality factor 3. F. Wavelength (microns) of wrinkled shapes as function of (*h*_0_/*R*).

The first term represents the stretching energy whereas the second term represents the bending energy. Minimizing the buckling energy with respect to the width of the bending strip *a* yields 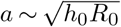. This leads to an estimate for the energy of the buckled actin shell:

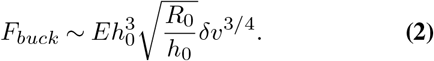

The maximal curvature (located at the edge of the bending strip) reads,

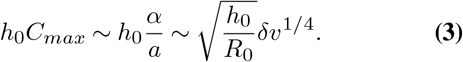

The maximal curvature therefore depends on the liposome radius and sub-linearly on the volume change *δυ*. In order to check this scaling, we compare the experimentally obtained shapes for the buckled shells with Eq. 3 in Fig. 2C. The shapes of the actin shells follow the predicted scaling of the maximal radius of curvature. The only unknown in Eq. 3 is a geometrical constant of order one. In fact, a constant of unity captures well our data (Fig. 2C). Note that this relation is valid for *h*_0_/*R*_0_ ≪ 1 and hence it is expected to break down for thick actin shells, as observed in Fig. 2D.

### Wrinkled shapes

Our energy estimates can explain the generic shape of *buckling* for thin actin shells. Thick actin shells display *wrinkling*. Such a wrinkled shape is reminiscent of what is observed in curved multilayered surfaces, which are common to a wide range of systems and processes such as embryogenesis, tissue differentiation and structure formation in heterogeneous thin films or on planetary surfaces, or simply, a bilayer system consisting of a stiff thin film on a spherically curved soft substrate (22). Here, the stiff thin film is the membrane. Its bending rigidity *κ* is of the order of a few *kT*, close to thermal energy (18). The curved soft substrate is the actin network, characterized by an elastic Young modulus *E* of 10^3^-10^4^ Pa (23). One important feature of wrinkled shapes of such bi-layered substrates is that their wave length λ only depends on the bending modulus of the stiff thin film (here the membrane) and the elastic Young modulus of the soft substrate (here the actin network) and is therefore independent of the thickness of the actin layer (24):

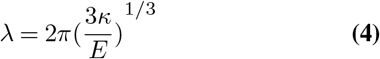

Estimates of the wrinkling wavelength of the membrane in our experiments for different sizes of liposomes and network thicknesses reveal that indeed, we find no dependence as a function of the normalized parameter *h*_0_/*R*_0_ (Fig. 2F).

There is a striking difference between the morphologies of the thinnest and the thickest actin shells categorized in Fig. 1D. For buckled shapes, the membrane deformation perfectly follows the one of the thin actin shell (Fig. 1C, left). In contrast, for thick actin shells, whereas the membrane clearly wrinkles, the outer actin shell boundary is undeformed as it appears perfectly spherical, and unaffected by membrane wrinkling (Fig. S2). This observation confirms that for wrinkled shapes, the elastic deformation does not penetrate the actin layer, but only on a depth on the order of the wavelength (25, 26). Intermediate actin shell thicknesses are discussed in detail below. Taking a *κ* value of about 10 *kT*, and an elastic Young modulus of about 10^3^ Pa (23), we find from Eq. 4 a λ on the order of 350 nm, which is in the lower range of experimentally estimated wavelengths (Fig. 2F), but in qualitative agreement with the occurrence of wrinkling happening for actin thicknesses above a few hundreds of nanometers (Fig. 1D).

### Buckling to wrinkling transition

In addition to the elastic properties of the actin layer and the membrane, the stability of the vesicle depends on the relative volume change *δυ* and on three length scales: the radius *R*_0_, the actin thickness *h*_0_ and the wrinkling wavelength, λ. After the osmotic shock, the relative volume change is fixed and the state of the vesicles depends on the two reduced variables λ/*h*_0_ and *h*_0_/*R*_0_. Large values of λ/*h*_0_ lead to a buckling instability and large values of *h*_0_/*R*_0_ lead to a wrinkling instability.

A detailed stability diagram showing the 3 possible states of the vesicle, non deformed, wrinkled and buckled is given in the Supplementary Materials using linear stability analysis. A remarkable property is that the transition line between buckled and wrinkled states is independent of the reduced volume *δυ* and can be written as a relation between *h*_0_ and *R*_0_ depending on the wrinkling wavelength λ

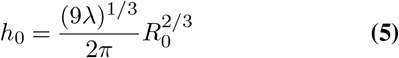

In order to test the mechanical model, in Fig.3, we plot all the experimental results corresponding to different values of *δυ* and of the 3 length scales in a plane (*h*_0_,*R*_0_). We observe a well defined separation curve between buckled and wrinkled states that we fit with Eq.5 by adjusting the value of λ. We obtain a value λ ≈ 2 *μm*, in reasonable agreement with the directly measured wavelength. The prefactors that we ignored in the scaling analysis are therefore of order 1.

**Fig. 3.**
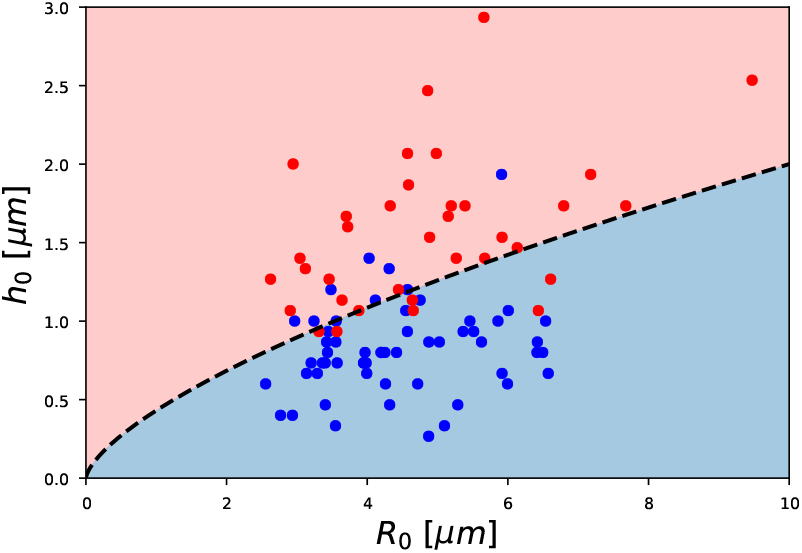
Buckled (blue dots) and wrinkled (red dots) shapes plotted in a (*h*_0_,*R*_0_) diagram. The dashed line is the best fit of data points with Eq. 5 and separates regions where buckling (blue) or wrinkling (pink) occurs.

## Conclusions

By combining experimental observation and theoretical analysis, we show here that the morphological changes of a liposome covered with an actin shell in response to an osmotic shock can be accurately controlled by monitoring the time of incubation and hence the thickness of the actin shell. We predict, (i) under which condition buckling or wrinkling occurs and (ii) the magnitude and spatial extend of deformation as function of the thickness of the actin shell and the osmotic compression. Comparing the experimentally observed shapes with the theoretical predictions for the transition from buckling to wrinkling demonstrates how actin shell mechanics controls the mode of deformation.

As shown in Fig. 1C, the surface of a wrinkled liposome displays small spatial undulations around the average radius *R*, which remain unchanged over time and are therefore clearly distinct from thermal fluctuations. The wrinkling undulation occurs at wavelengths much smaller than the radius of the liposome, an observation that is similar to mimics of actomyosin cortices (19). Local bending is favored into a buckling shape whereas wrinkling implies deformations all along the actin shell. Because the bending energy is too costly for thick actin shells, buckling is less stable than wrinkling. As a consequence, wrinkled shapes are only found if the actin thickness is larger than the wavelength. Experimentally, wrinkling of acto-myosin shells within incompressible oil droplets happens after a critical time that depends on the presence of a compressive stress energy term, due to acto-myosin contraction (19). In our system, the change in volume due to the osmotic shock may create compressive stresses of the thick actin layer in the absence of myosin, a phenomenon that would slow down localized shell buckling, and therefore favor wrinkling we observe here for thick actin shells. Furthermore, a small change in volume leads to wrinkling whereas a higher change leads to buckling (Fig. 1C), in agreement with the reasoning that bending into a buckled shape for thicker actin shells is more costly in energy than thinner actin shells, and therefore, smaller liposomes deform by wrinkling. Note that the elastic Young modulus of the actin network has no effect on the shape of the buckling deformation, and only weakly affects wrinkled shapes (1/3 exponent, Eq.4).

Our results are in line with various observations of localized deformation in constrained growth in mechanical systems, such as the wrinkling to fold transition observed in various supported elastic media (8, 27–29), the shaping of the brain (30), and the buckling transition in virus shells (31, 32) and pollen grains (33). One major advantage of our reconstituted system is our ability to finely tune the structural parameters of the actin shell (mostly its thickness here) so as to explore the phase diagram of possible shape transitions. Tuning the incubation time of the branched actin network or shell thickness and the amount of osmotic compression of the shell, directly controls the intrinsic mechanical response and applied force that sets the deformation.

## Acknowledgements

This work was supported by the French Agence Nationale pour la Recherche (ANR), grant ANR-14-CE090006 and ANR-12-BSV5001401, and by the Fondation pour la Recherche Médicale (FRM), grant DEQ20120323737. Thanks to the Bettencourt Schueller Foundation long term partnership, this work was partly supported by CRI Research Fellowship to Remy Kusters. Cécile Sykes would like to thank the Gordon Research Conferences, in particular the conference on Soft Condensed Matter Physics that was held on August 11-16, 2019 (Colby-Sawyer College, New London, NH, USA; chairs: Lisa Manning and Christoph Schmidt), for promoting extremely lively and insightful discussions on soft matter physics and biological systems. Special thanks go to Andrej Kosmrlj for presenting an update on wrinkling phenomena and for his didactic training on the wrinkling literature. Thanks also to Robijn Bruinsma for checking the theoretical interpretation of buckling and wrinkling, and Jacques Prost and Catherine Quilliet for shrewd discussions.

## Supplementary Note 1: Actin shells control buckling and wrinkling of biomembranes

### Linear stability analysis

We discuss in this section the stability of a spherical actin shell around a vesicle at the scaling level. We use the notations of the main text, the liposome has a radius *R*_0_ and the lipid membrane a bending modulus *κ*. The actin shell, which we consider incompressible for simplicity, has a thickness *h*_0_ and a Young modulus *E*. The deflation of the membrane creates a relative decrease of the vesicle volume −*δυ* and induces a negative tension −*γ* of the actin layer with 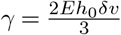.

We first ignore the curvature of the surface and discuss the stability of a flat membrane covered with an elastic layer of thickness *h*_0_ under a negative tension −*γ*. We consider an undulation of the membrane *u*cos*qx*. This deformation creates an elastic stress in the elastic layer and costs elastic energy. In the limit where the wave vector is small, *qh*_0_ ≪ 1 the elastic energy per projected area corresponds to the bending of a thin elastic sheet 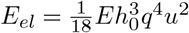. In the opposite limit where the elastic layer is thick *qh*_0_ ≫ 1, the elastic deformation only penetrates over a length 1/*q* inside the elastic layer and the elastic energy per projected area is 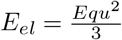. As the relevant length scales are well separated, it is sufficient for our purpose to extrapolate the elastic energy per projected area between these two limits

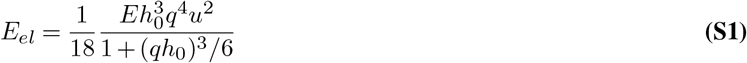

The total energy *F* per projected area is the sum of the bending energy of the membrane, the elastic energy per projected area *E_el_*, and the negative tension (−*γ*). The stability of the layer is best discussed in terms of its effective tension. We define the effective tension by writing the total energy as 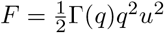. The surface of the layer becomes unstable with respect to undulations when the effective tension vanishes. The effective tension is given by

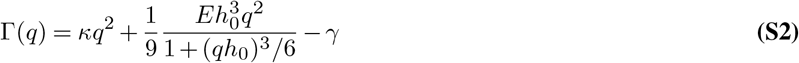

If the thickness of the elastic layer *h*_0_ is larger than λ, the tension is a non monotonic function of the wave vector *q* with two minima, a minimum at zero wave vector corresponding to buckling and a minimum at finite wavevector corresponding to wrinkling. The wrinkling wave vector is

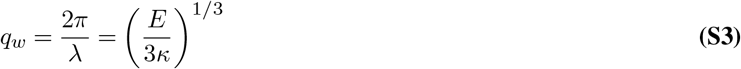

If the thickness *h*_0_ of the elastic layer is smaller than the wave length λ the wrinkling minimum disappears and the only instability is buckling. At the scaling level, for a spherical vesicle of finite radius, one can still study the instability from the tension Γ given by Eq. S2. However, the smallest wave vector where buckling can occur is *q* = 2*π*/*R*_0_.

To fully describe the transition between buckling and wrinkling, we build a diagram of states of the liposomes looking at the instability with respect to both of buckling and wrinkling for which Γ = 0. We obtain the two lines of Fig. S3 that limit the domains for buckling and wrinkling: using the expression of *γ*, we find that the vesicle is unstable with respect to buckling if *h*_0_/*R*_0_ ≤ (*δυ*)^1/2^ and it is unstable with respect to wrinkling if λ/*h*_0_ ≤ *δυ*. The relative volume change *δυ* is fixed and we describe the state of the vesicle in the plane of the two dimensionless variables (λ/*h*_0_,*h*_0_/*R*_0_) (Fig. S3).

The transition between buckling and wrinkling occurs when the effective tension Γ is equal in the buckled and wrinkled states. This transition is independent of *δυ* and occurs if

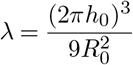

We compare this equation to the experimental results in the main text.

### Supplementary Figures

**Fig. S1.**
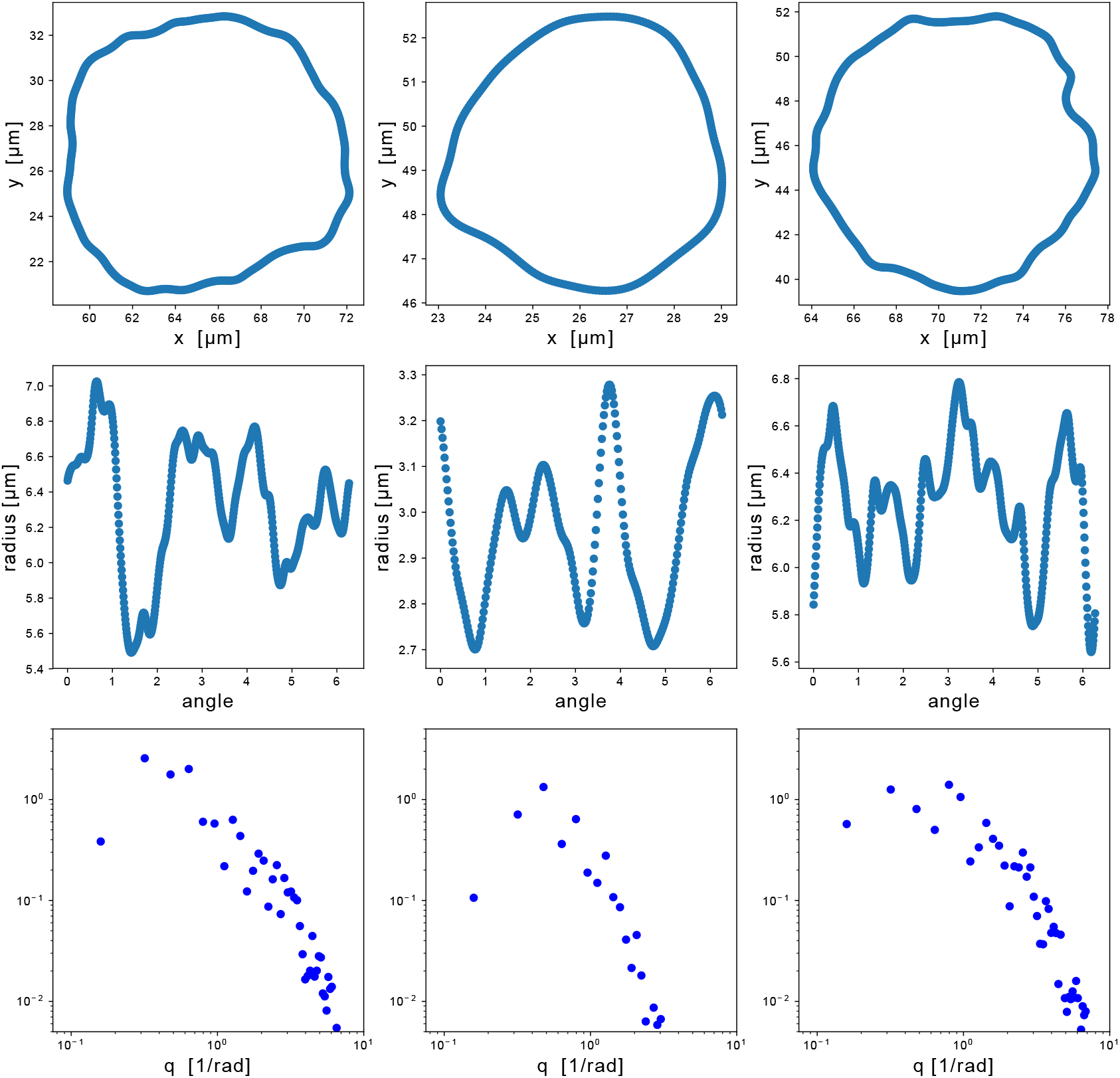
Three examples of wrinkling (top row) with their corresponding radius as function of the angle (middle row) and their Fourier spectrum (bottom row). The maximal value of the Fourier spectrum is selected as characteristic wavelength.

**Fig. S2.**
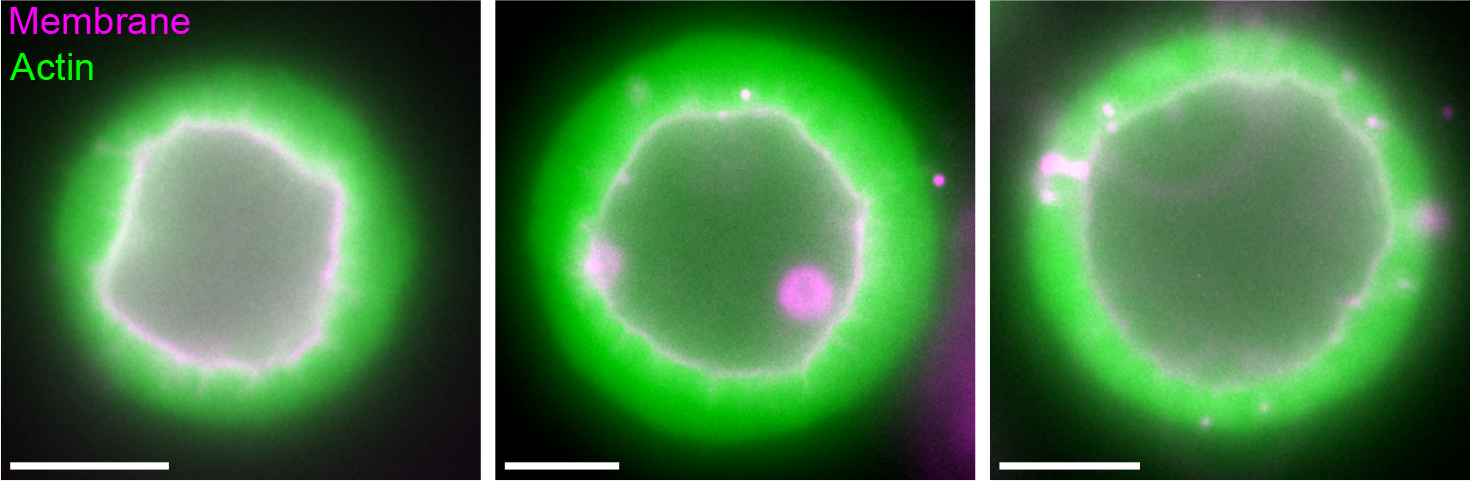
Three examples of membrane (magenta) and actin (green) in the case of wrinkled shapes. Note that the outer contour of the actin network is spherical and un-deformed whereas the membrane wrinkles. Scale bars: 5 *μ*m.

**Fig. S3.**
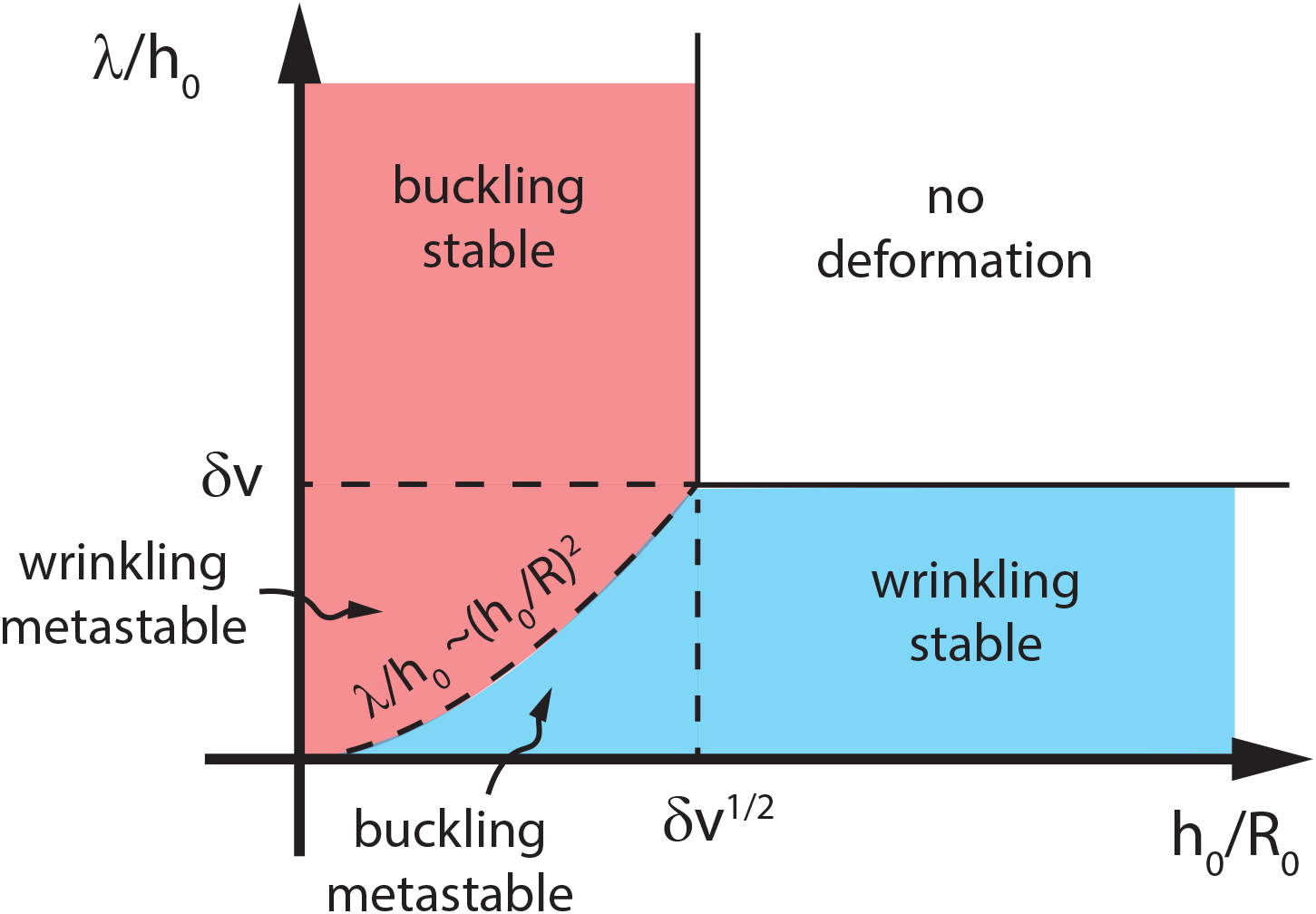
Stability diagram of a spherical vesicle with an actin shell undergoing both wrinkling and buckling. The dotted line represents the equality of tension *γ* for both buckling and wrinkling.

### Supplementary Movies

**VideoS1**. Naked liposome when preparation evaporates over time. Equatorial plane observed by epifluorescence as a function of time. Bar, 5 *μ*m.

**VideoS2**. Naked liposome under osmotic shock. Equatorial plane observed by epifluorescence as a function of time. Time indicated is the time after the osmotic shock. Bar, 5 *μ*m.

## Footnotes

1 The python/numpy notebooks containing the analysis are available on Github: *https://github.com/remykusters* and are based on the SciPy cookbook, *https://scipy-cookbook.readthedocs.io/*

